# Functional transcriptional signatures for tumor-type-agnostic phenotype prediction

**DOI:** 10.1101/2023.04.12.536595

**Authors:** Corey Weistuch, Kevin A. Murgas, Jiening Zhu, Larry Norton, Ken A. Dill, Allen R. Tannenbaum, Joseph O. Deasy

## Abstract

Cancer transcriptional patterns exhibit both shared and unique features across diverse cancer types, but whether these patterns are sufficient to characterize the full breadth of tumor phenotype heterogeneity remains an open question. We hypothesized that cancer transcriptional diversity mirrors patterns in normal tissues optimized for distinct functional tasks. Starting with normal tissue transcriptomic profiles, we use non-negative matrix factorization to derive six distinct transcriptomic phenotypes, called archetypes, which combine to describe both normal tissue patterns and variations across a broad spectrum of malignancies. We show that differential enrichment of these signatures correlates with key tumor characteristics, including overall patient survival and drug sensitivity, independent of clinically actionable DNA alterations. Additionally, we show that in HR+/HER2-breast cancers, metastatic tumors adopt transcriptomic signatures consistent with the invaded tissue. Broadly, our findings suggest that cancer often arrogates normal tissue transcriptomic characteristics as a component of both malignant progression and drug response. This quantitative framework provides a strategy for connecting the diversity of cancer phenotypes and could potentially help manage individual patients.

## INTRODUCTION

Recent evidence has emerged that large-scale patterns of transcriptional activity often follow similar rules across different cancers [1–6]. For example, fast growing tumors typically exhibit the “Warburg effect” (metabolic reprogramming to selectively upregulate glycolysis), while slower growing tumors (as often observed following senescence-inducing chemotherapies) commonly upregulate oxidative metabolism [7]. These and other realizations have fueled significant recent interest in targeting cancer phenotypes characterized by systems-level properties rather than considering only a limited set of “actionable” genetic mutations [8, 9]. However, fundamental questions remain unanswered: What do such large-scale transcriptomic patterns represent? How do we identify these patterns using molecular data? Most urgently, how can we exploit the identified patterns to guide therapy?

Due to the vast diversity of cancer types and the numerous potential patterns that can be identified within them, there is currently no universally accepted standard for classifying transcriptomic phenotypes in cancer [2–4]. Unlike tumors, however, normal tissues typically have well-separated gene expression profiles and defined functional roles. Furthermore, many of the transcriptional differences observed across cancers mimic patterns of variability observed in normal tissues, making them an ideal system for contextualizing observed patterns of intertumor and intratumor heterogeneity [10, 11]. Leveraging these links could therefore unveil new and general connections between the welldefined functional roles of normal tissues and recurring patterns of tumor heterogeneity.

Here, we use a statistical decomposition method to isolate and interpret transcriptomic motifs utilized both by normal tissues and tumors, referring to these signatures as *normal tissue archetypes* [12]. By restricting our analysis to biological pathways utilized by all tissues, such as glycolysis and DNA repair, the derived archetypes are mapped to unique tasks performed by normal tissues as well as distinct cancer hallmarks [2, 9, 13]. Moreover, our employment of a common reference has the potential to address existing challenges in comparing transcriptional diversity patterns across multiple cancer types and datasets [2–4]. Specifically, the approach calculates the mix of archetypes within a given cancer cell or bulk tumor in terms of fixed normal tissue archetype signatures, providing a new perspective and a flexible quantitative approach to classify the different possible types of tumor behavior, match them to their most effective treatments, and interpret how they evolve in response to various therapies and through the accumulation of genetic alterations.

In assessing the prognostic potential of normal tissue archetypes identified through our method, we employed a structured case-study approach, utilizing publicly available data from two pan-cancer datasets [14–17] and three datasets focusing on therapy-induced and metastatic adaptation in specific malignancies [18–20]. Our primary aim was to gauge the predictive capability of inferred archetype mixtures in determining the responsiveness of cancer cell lines to distinct chemotherapies, agnostic to mutational status, copy number alterations, and cancer type. Given the recognized divergence in behavior between cancer cell lines and clinical samples, we also scrutinized their ability to predict overall patient survival, differentiate between prognostic cancer subtypes, and anticipate adaptive responses from bulk tumor transcriptomes.

In a broader context, this proof-of-concept study highlights the prognostic significance derived from understanding why tumors commonly adopt normal tissue transcriptional programs distinct from their lineages of origin. The ability to identify and link these programs to drug sensitivities, site-specific metastases, and DNA alterations may pave the way for more tumor-type agnostic personalized cancer therapies.

## METHODS

### Ethical compliance

All analyses used publicly available, de-identified data.

### Public datasets

#### GTEx

We utilized the normal tissue samples from the Genotype-Tissue Expression (GTEx) project to cover the breadth of gene expression space and to provide an enhanced signal for resolving transcriptomic archetypes of both normal and cancerous tissues [21]. GTEx (version 8) gene-level transcripts per million (TPM)-normalized expression data were downloaded from the GTEx Portal [21]. The dataset, consisting of 54 distinct tissues, is one of the most comprehensive resources for studying tissue-specific gene expression. GTEx was established to characterize the tissue-specific determinants of human traits and diseases and provides expression levels for about 44 thousand genes. We focus here on commonly enriched pathways in cancer by utilizing 780 genes from five key cancer-related pathways from the Molecular Signature Database (MSigDB): apoptosis, DNA repair, glycolysis, hypoxia, and oxidative phosphorylation [22, 23]. To go beyond current classifications, we filtered out lineage-specific genes already used for cancer classifications, enabling comparisons across cancer types [24]. The above five gene sets (“pathways”) were chosen as a precaution against overfitting the 54 median-averaged bulk transcriptomes available in GTEx and because they play particularly frequent roles in cancer [8]). This ensures that our analysis is of relevance to multiple types of cancer. The remaining hallmark pathways, not encompassed by the previously mentioned patterns, included either specific signaling pathways or covered functions that are not ubiquitously expressed across tissues. Finally, to normalize the dataset, we divided the expression of each gene by its standard deviation across all tissues.

#### Cancer Cell Line Encyclopedia, CCLE

Gene-level TPM-normalized expression from the CCLE (*N* = 1405) along with matched drug sensitivities (*N* = 469), mutations (*N* = 1250), and copy number alterations (*N* = 1387 were downloaded from the public Dependency Map (DepMap) portal (version 22Q2) [14]. To reduce false positives, only TCGA hotspot mutations present in at least five TCGA samples (as reported in CCLE) and ten CCLE samples were retained. Mutation types were not further stratified. Due to the lack of a similar reference for pan-cancer copy number analysis, all whole genome-level copy number alterations reported in CCLE, aside from those on the sex chromosomes, were used. The upper bound for gene-level copy number alterations was set to four in order to remove potential correlational biases from high copy number states. Spearman rank correlations were then computed between the copy number state of each gene and each computed normal archetype score. Finally, region-level correlations were computed by resegmenting the genome into 20000 bins with equal mappability, averaging gene-level correlations within each bin.

#### Gastrointestinal Stromal Tumor (GIST) therapy responses

Gene-level TPM-normalized expression data from imatinib-sensitive (*N* = 5) and imatinib-resistant (*N* = 5) GIST patients were downloaded from NCBI GEO accession code GSE155800 [18].

#### Longitudinal study of metastatic breast cancer

Gene-level TPM-normalized expression data from east Asian HR+/HER2metastatic breast cancer patients before and after treatment with palbociclib plus endocrine therapy (*N* = 23 matched pairs) were downloaded from NCBI GEO accession code GSE186901 [19].

#### The Cancer Genome Atlas, TCGA

Gene-level TPM-normalized expression data from the TCGA within the breast (BRCA, *N* = 1111), colon (COAD, *N* = 481), and pancreatic (PAAD, *N* = 178) cancer cohorts were downloaded via the TCGAbiolinks R package [15–17, 25]. Samples were restricted to primary tumors.

#### Study of site-specific adaptation of metastatic breast cancer

Raw sequencing data from the basal (triple negative) metastatic breast cancer patients used in this study (*N* = 7; *N* = 5 caucasian, *N* = 2 african) are available for controlled access from dbGaP accession code phs000676.v2.01 [20]. Gene-level quantile and TPM-normalized expression data were downloaded from GitHub https://github.com/FaribaRoshanzamir/Metastatic-TNBC/tree/main/data and were additionally processed as documented in [26]. In summary, primary tumors from each patient were used alongside their matched distant metastases to brain (*N* = 7), lung (*N* = 6), liver (*N* = 5), lymph node (*N* = 2), adrenal gland (*N* = 2), and skin (*N* = 2). The samples were then batch-corrected against normal tissue samples from GTEx and primary tumor samples from TCGA corresponding to each metastasis destination. Median expression levels of each metastasis site, normal tissue, and primary tumor were then provided. Due to the lack a of matched normal tissue in GTEx, the sample derived from lymph node metastasis was excluded from our analysis.

### N-NMF archetype analysis

Biological data often embody the amalgamation of interconnected constituent parts. Our goal is to isolate the parts mathematically, revealing key structures and hidden patterns. This is commonly accomplished through approximate low-rank matrix and tensor factorizations, including principal component analysis (PCA) and non-negative matrix factorization (NMF). Notably, NMF has played a significant role in recent biological studies, contributing to dimension reduction, discrimination, and clustering [3, 4, 27]. When the factor weights are normalized (N-NMF), they encode the relative contribution of each part in contributing to the whole [28]. Here we detail our implementation of this factor or “archetype” discovery procedure.

#### Choosing the optimal number of archetypes

Application of NMF requires predefining the number of archetypes. The optimal number of archetypes was chosen using the profile log-likelihood method [29]. This approach models the unknown number of archetypes as a latent variable that can be directly optimized over. This method was selected due to its simplicity and superior performance compared to alternatives [30]. When training the archetype model on normal tissue transcriptomic data (GTEx), the minimum number of factors was set to *k* = 2 (due to the normalization constraint), with the maximum set at *k* = 30. The optimal number of archetypes, determined by maximizing the resulting profile log-likelihood, was determined to be *k* = 6 (cf. SI Fig. 1).

#### Archetype discovery and sample projection

Normalized nonnegative matrix factorization (N-NMF) was employed to establish a low-dimensional representation of gene expression profile variability across normal tissues (GTEx). This general approach seeks to approximate an *N × M* data matrix *V*, representing *N* gene expression values of *M* samples, as the product of two low-rank matrices *W* and *H*: *V* ≈ *WH*, where *W* is an *N × k* matrix representing coefficients of each gene’s contribution to the *k* = 6 archetypes, and *H* is a *k × M* matrix representing the weights of each archetype needed to approximate each gene expression sample (cf. Fig. 1A). In our implementation, these matrices are found by minimizing the Frobenius norm distance between the data matrix *V* and its approximation [28]. The optimal solution is obtained through 1000 iterations of the classical Seung-Lee algorithm [31]. Prior to this, the *W* and *H* matrices are randomly initialized using a uniform distribution. To avoid degeneracy in the model fitting and ensure that the coefficients of each archetype score sum to 1, the *H* matrix is column-normalized after each iteration [28]. Subsequently, when calculating archetype scores for new samples, we follow the same procedure but fix the *W* matrix to its learned values.

**FIG. 1:**
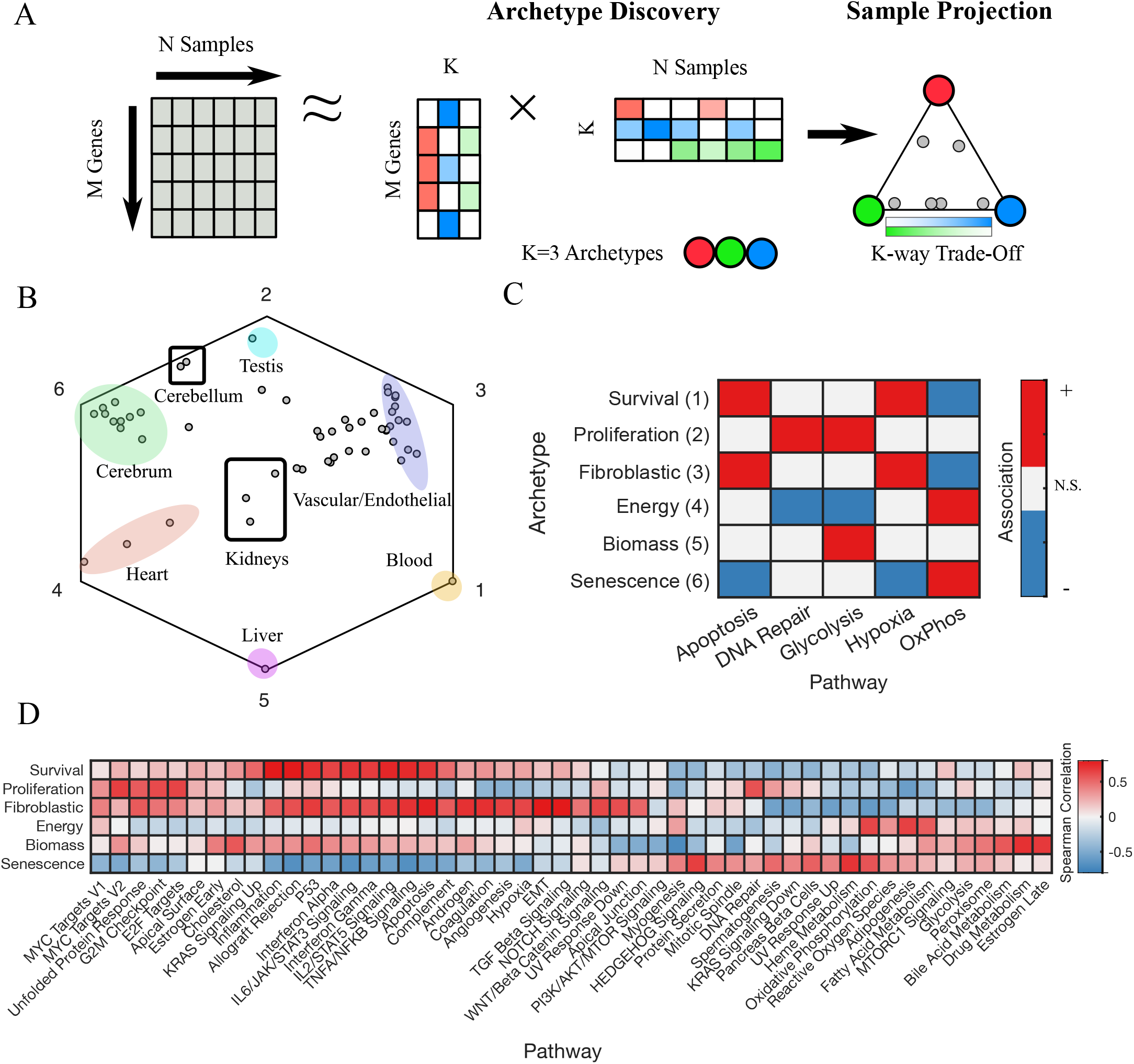
Normal tissue archetypes as a paradigm to interpret cellular behaviors. (A). Flowchart of the analysis pipeline (see Methods: N-NMF archetype analysis). (B). Projection of median-averaged normal tissue transcriptomes onto the six archetypes discovered in GTEx (*N* = 54 distinct tissues). The coordinates of each tissue correspond to their degree of similarity to each archetype (the vertices). Tissue groups enriched for a single archetype are marked by ovals, whereas those achieving a balance of multiple archetypes are marked by squares. (C). Heatmap of the significant associations between individual archetypes and the six pathways on which they were trained (Spearman rank permutation test, Benjamini-Hochberg *p <* 0.05). (D). Heatmap of the Spearman correlations between individual archetypes and the average expression levels of each MSigDB Hallmark gene group.

### Statistical Analysis

Statistical analyses were performed using MATLAB 2020b. Associations between archetype scores and all other variables were determined using paired or unpaired Spearman rank correlation t-tests unless otherwise indicated. The Benjamini-Hochberg procedure for multiple comparisons correction was applied when appropriate and as indicated [32]. Associations with *p <* 0.05 were considered significant unless otherwise indicated. Kaplan-Meier survival analyses were performed using MatSurv [33].

## Data and code availability

The source data and MATLAB code that support the findings of this study are available in GitHub with the identifier https://github.com/Corey651/Cancer_Archetypes.

## RESULTS

### Normal tissue diversity signatures mimic oncogenic transcriptional programs

Normalized Nonnegative Matrix Factorization or N-NMF (cf. Methods and Fig. 1A, [28]) was used to identify the most representative archetypes of the normal tissue transcriptomes in the Gene-Tissue Expression Project (GTEx, cf. Methods). The method determined that the data was best represented by six archetypes (cf. SI Fig. 1, Methods), each associated with distinct normal tissue types (cf. Fig. 1B) and biological pathways (cf. Fig. 1C,D). Importantly, each normal tissue was characterized by a mixture of archetypes, although one is typically dominant. Related tissues, such as the constituents of the cerebrum, were seen to have more similar archetype scores, in accordance with expectations. Crucially, the presence of single-tissue archetypes (e.g., testis and liver) and distinct clustering into interpretable tissue classes suggest that our analysis avoids bias towards overrepresented tissue classes.

Although the initial analysis focused on select ubiquitously utilized pathways, Figure 1D illustrates notable Spearman rank correlations among numerous additional Hallmark gene pathways sourced from MSigDB and the individual archetype scores [22, 23]. Archetype 1 exhibited enrichment for immune pathways, including interferons, TNFA/NFKB signaling, and apoptosis, suggestive of an anti-apoptotic, immune-evasive phenotype termed “Survival” [34]. Archetype 2 demonstrated enrichment for cell division pathways such as the G2/M checkpoint, MYC targets, and DNA repair, associated with “Proliferation” [35]. Archetype 3, sharing some immune-related pathways with Archetype 1, also displayed enrichment for vascularization (angiogenesis), cell adhesion (apical junction), and cell signaling pathways (NOTCH, TGFB), collectively associated with “Fibroblastic” activity [36]. Archetype 4 was enriched for catabolic or “Energy” metabolism pathways. Archetype 5 was enriched for hormonal (estrogen, cholesterol), drug metabolism, and glycolysis pathways, collectively reflecting known liver functions summarized as “Biomass”. Archetype 6 showed enrichment for heme metabolism, oxidative phosphorylation, and HEDGEHOG signaling reflective of a differentiated phenotype. Additionally, archetype 6 exhibited broad negative associations with immune-related pathways, contrasting with archetype 1, reminiscent of “Senescent” cells [37].

Of note, the scores recapitulate known functional trade-offs across normal tissues, such as the elevated oxidative requirements of the heart and cerebrum (“Energy” and “Senescence”), the requirement for glycogen production in the liver (“Biomass”), and the protection, regulation, and manipulation of genetic material in the testis (“Proliferation”). Notably, certain tissues encompass multiple archetypes, thereby striking a balance between various functions. For instance, the cerebellum, akin to the testis [38, 39], demonstrates resilience to aging and accumulated DNA damage, while also exhibiting oxygen-demanding characteristics similar to the cerebrum [40]. The kidneys, expressing approximately 70% of the genes in the human body [41], assume intermediate archetype values, reflecting their versatile functional profile.

Crucially, the normal tissue archetype-association enrichment patterns also resemble groups of pathways commonly co-expressed in many different types of cancer [2, 8, 13]. Specifically, several of the archetypes closely correspond to those previously identified in tumors: Survival → Immune interaction, Proliferation → Cell division, and Fibroblastic → Invasion/tissue remodeling [13]. Energy and Biomass, on the other hand, were somewhat similar to combinations of these existing signatures, whereas Senescence bore no such resemblance. Of note, several chemotherapies induce a treatment-resistant, senescence-like states in tumors, characterized by immunogenic and metabolically-active features (termed SASP) akin to those of the Senescence archetype [37].

### Non-canonical tissue signatures are enriched in cancer cell lines

To determine how well normal tissue archetypes capture shared expression patterns across cancers compared to their respective lineages of origin, we analyzed their distribution across the Cancer Cell Line Encyclopedia (*N* = 1405, [14, 42], cf. Methods). Consistent with typical tumor expression patterns, the cancer cell lines were predominantly enriched for the Proliferation and Senescence archetypes [37] with minor variation across lineages (cf. Fig. 2A). However, while certain cancer types exhibited preferences for specific archetypes, these preferences (with the exception of the liver) did not mirror those of their respective normal tissue lineages (cf. Figs. 2B,C).

**FIG. 2:**
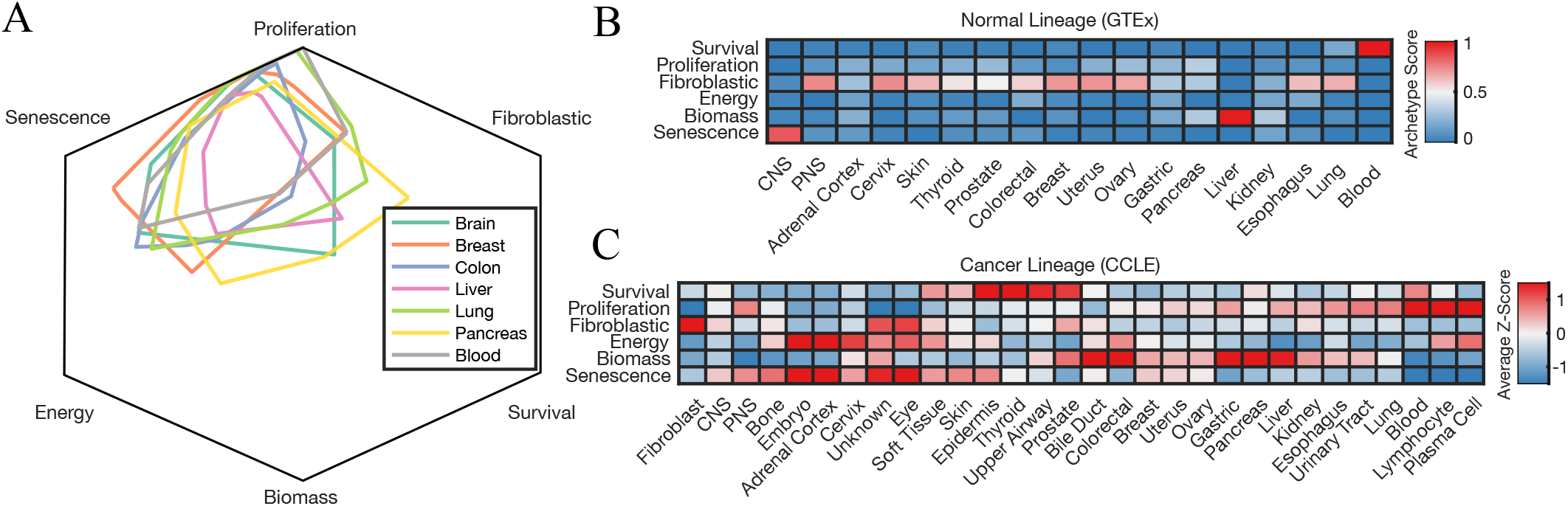
Variation in normal archetype enrichment across cancer cell line lineages. (A). Ranges of archetype values in CCLE, stratified by lineage. Archetype ranges were calculated using convex hull estimation. (B). Heatmap of the archetype patterns from normal tissues (GTEx) with corresponding cancer cell line lineages, displaying the first matched tissue type for each lineage. (C). Heatmap of the average archetype Z-scores across tumor lineages (CCLE), computed by averaging expression within each lineage and calculating Z-scores relative to the overall data. Z-scores were used to correct for the average archetype trend observed in cancer cell lines (A).

### Normal tissue signature enrichment influences pan-cancer drug responses

We next investigated the ability of normal tissue archetypes to predict tumor drug sensitivities across a variety of cancers, conditions, and data types. We hypothesized that their connections to frequently dysregulated cancer pathways, along with their similar distributions within different cancer types, could offer a strategy for addressing common challenges faced by existing predictive models related to their lack of interpretability, difficulty generalizing to new datasets, and their limited ability to monitor changes in drug sensitivities over time [43].

As illustrated in Figure 3, transcriptomic signatures derived from normal tissues demonstrated the capability to predict drug sensitivities across diverse cancer types, identifying both cell lines and patients sensitive to treatment. Notably, cancer cell line drug sensitivities (quantified by activity area, cf. [14]) of 21/24 anti-cancer drugs profiled by CCLE exhibited a significant association with at least one archetype, following pooling across cancer types and multiple comparisons correction (cf. Fig. 3A). Moreover, these associations aligned with the pathway-specific characteristics of the archetypes (cf. Fig. 1D). For instance, archetypes enriched for apoptosis-related (Survival), DNA repair (Proliferation), and hormonal pathways (Biomass) were associated with sensitivities to *XIAP* anti-apoptosis inhibitors (LBW242), topoisomerase inhibitors (irinotecan and topotecan), and *EGFR* inhibitors (lapatinib and erlotinib), respectively.

**FIG. 3:**
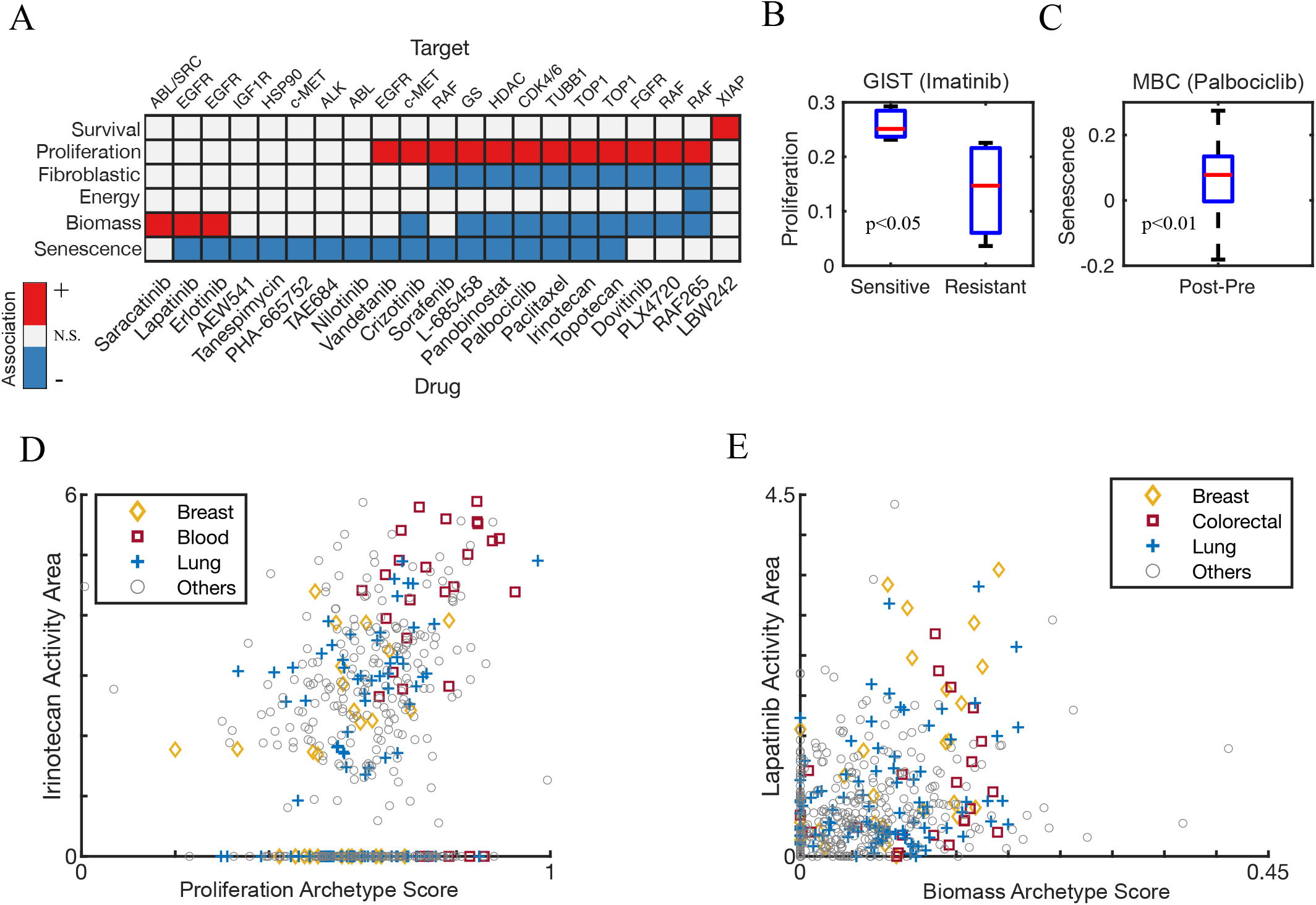
Associations between normal archetype scores and chemotherapy sensitivities across a range of cancer types. (A). Heatmap of the significant associations between individual archetypes and the cancer cell line activity areas of each measured drug (Spearman rank permutation test, Benjamini-Hochberg *p <* 0.05). (B). Boxplots comparing the Proliferation archetype scores in imatinib-sensitive and resistant GIST patient samples (t-test, *N* = 10, [18]). (C). Boxplot of the change in Senescence archetype scores in HR+/HER2metastatic breast cancer samples (MBC) before and after treatment with palbociclib (paired t-test, *N* = 23, [19]). (D). Scatterplot showing the Proliferation archetype score vs irinotecan activity area, stratified by cancer lineage. *R*^2^ values for lineage-specific linear regressions were 0.2199, 0.2152, and 0.1689 for breast, blood, and lung, respectively. (E). Scatterplot showing the Biomass archetype score vs lapatinib activity area, stratified by cancer lineage. *R*^2^ values for lineage-specific linear regressions were 0.2558, 0.0465, and 0.0794 for breast, colorectal, and lung, respectively. Cancer lineages shown in D and E were manually selected based on their total number of samples and average archetype expression levels. Cell lines with 0 activity area, indicating incomplete information, were excluded from analysis. To enhance visibility of all data points, regression lines were excluded.

Proliferation archetype scores were significantly reduced in gastrointestinal stromal tumor (GIST) patients resistant to imatinib (cf. Fig. 3B). Furthermore, treatment with palbociclib, known for eliciting therapy-induced tumor senescence as a resistance mechanism in various cancer types [44], was associated with significant increases in Senescence archetype scores in HR+/HER2metastatic breast cancer samples compared to their pre-treatment state (cf. Fig. 3C). Thus, archetype scores can also provide prognostic information about patients and how their responses to therapy change over time.

Even when stratified by cancer type, the Proliferation and Biomass archetypes were associated with increased sensitivity to irinotecan and lapatinib, respectively (cf. Figs. 3D,E). Although Survival was significantly associated with LBW242 sensitivity, it was rarely expressed in cancer cell lines and was therefore excluded from stratified analyses in this study. Lineage-specific *R*^2^ values ranged from 0.0465 to 0.2558, consistent with previous regression models on this dataset [14]. Of note, Figure 3E indicates a nonlinear threshold effect, whereby sensitivity to lapatinib in colorectal and breast cancer is observed solely in cell lines with Biomass archetype scores exceeding 0.15. Overall, our findings support the idea that archetypes reveal associations that withstand differences between cancer types.

### Normal tissue signature enrichment improves survival prediction and distinguishes site-specific metastases in breast cancer

Recognizing the promising capacity of normal tissue archetypes in predicting and monitoring treatment responses, we sought to assess their broader applicability in interpreting existing treatment guidelines and overall clinical outcomes. This encompassed three specific tasks. Firstly, conducting a comparative analysis between established treatment guidelines and those proposed by our archetypes, thereby illustrating the translational implications of our earlier discoveries. Secondly, evaluating the potential of these signatures for identifying vulnerable patients more effectively than existing stratification schemes. Lastly, characterizing archetype enrichment changes and potential archetype-predicted vulnerabilities associated with metastasis. To accomplish these tasks, we chose breast cancer as a pertinent case study due to its well-established transcriptional subtypes [45], diversity of potential metastatic sites [20, 26], and ample available data for study [15, 20].

To characterize subtype-specific archetype enrichment patterns, we used RNA-Seq data from breast cancer cell lines (CCLE, *N* = 63) and primary patient samples (TCGA, *N* = 1111, [15]). Both datasets encompassed standard breast cancer subtypes, with CCLE distinguishing two basal subtypes (A/B) and TCGA distinguishing two luminal subtypes (A/B) along with an additional normal subtype (cf. Figs. 4A,B). Perhaps owing to differences in purity between cell lines and clinical samples, absolute archetype scales differed between CCLE and TCGA. Furthermore, the correlation between breast cancer subtype and individual archetypes was limited, as evidenced by their substantial subtype-intrinsic variance, especially in TCGA (cf. Fig. 4B). Of note, however, both datasets revealed concordant archetype-enrichment trends consistent with existing subtype-specific breast cancer therapies. Basal tumors exhibited the highest median Proliferation scores, and HER2+ tumors showed the highest median Biomass scores across both datasets (cf. Figs. 4A,B). Consistent with these enrichment trends and our findings in Figure 3A, basal tumors are typically treated with DNA damaging agents, while HER2+ tumors are treated with a combination of trastuzumab and *EGFR* inhibitors including lapatinib. Furthermore, basal A, despite being described as having luminal-like properties [46], showed enrichment for Biomass, suggesting a closer resemblance to HER2+ and potential sensitivity to lapatinib and other EGFR inhibitors (cf. Fig. 4A). Luminal B (LumB), considered and intermediate phenotype between luminal A (LumA) and basal [47], also exhibited an intermediate median Proliferation score in TCGA (cf. Fig. 4B).

**FIG. 4:**
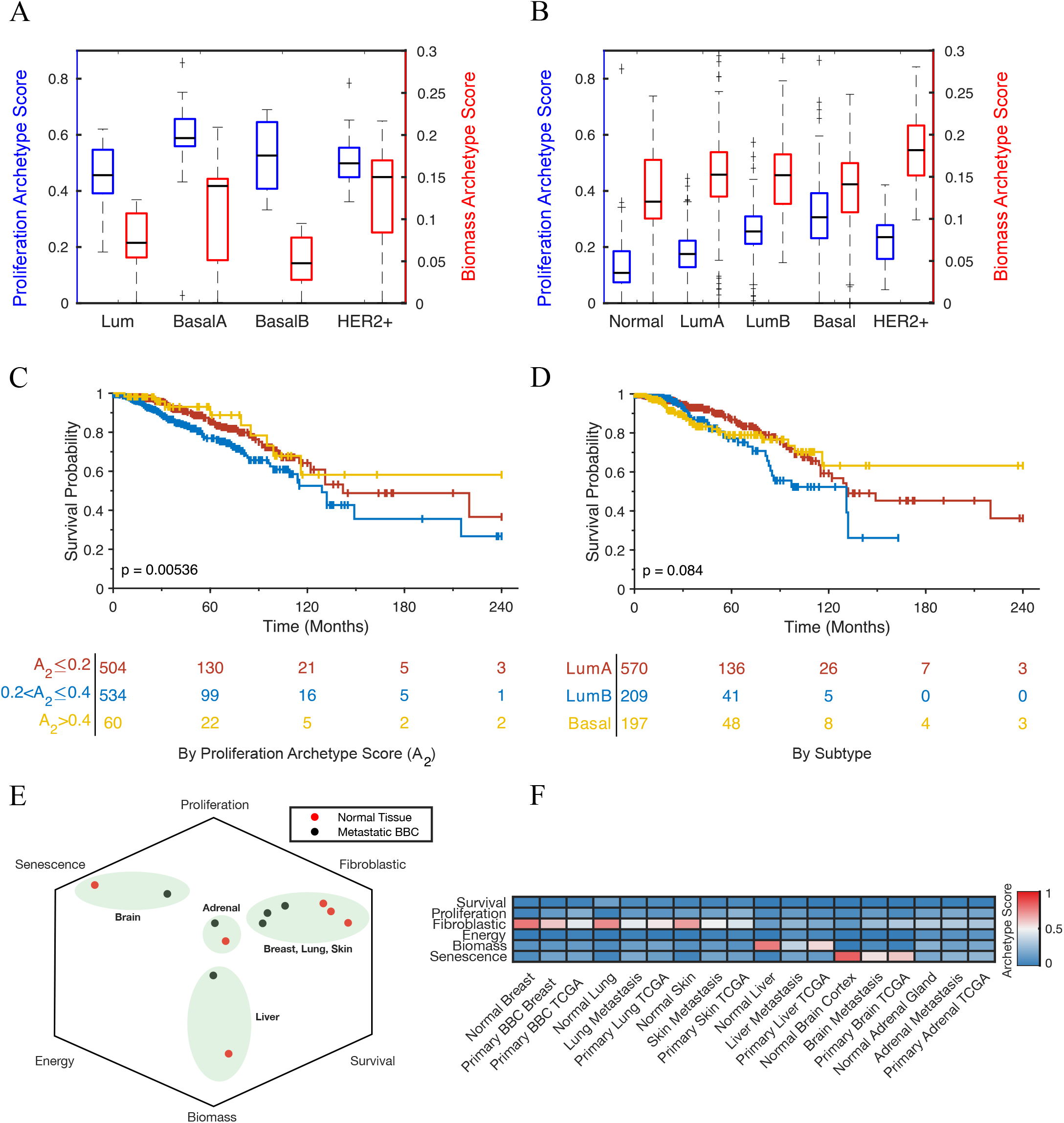
Associations between normal archetype scores and distinct breast cancer behaviors. (A). Boxplots of the Proliferation (blue) and Biomass (red) archetype scores across breast cancer cell lines, stratified by subtype. (B). Boxplots of the Proliferation (blue) and Biomass (red) archetype scores across primary breast tumors in TCGA, stratified by subtype. (C). Kaplan-Meier survival analysis, stratified by Proliferation archetype score. (D). Kaplan-Meier survival analysis, stratified by reported breast cancer subtype. Survival analyses utilized overall survival and RNA-Seq data provided by TCGA. Tables below indicate the number of surviving patients at each time point. Significance was determined using a log-rank test. (E). Projection of median-averaged site-specific basal breast metastases (BBC, black) and matched normal tissues from GTEx (orange) projected onto the six normal tissue archetypes. (F). Heatmap of the median archetype patterns from breast metastases, normal tissues, and primary tumors associated with each site of metastasis.

To evaluate whether archetype enrichment patterns can be used to predict overall survival in breast cancer, we conducted parallel Kaplan-Meier analyses on subgroups of patients in TCGA stratified by their archetype values (cf. Fig. 4C) and according to their reported breast cancer subtype (cf. Fig. 4D). Normal and HER2+ samples were excluded due to their limited representation and distinct treatment standards, respectively. Following our previous findings, the remaining patients were stratified by their Proliferation archetypes scores. Notably, patients with intermediate Proliferation archetype scores exhibited a significantly poorer prognosis compared to their counterparts (log-rank test, *p* = 0.00536). Furthermore, although patients with luminal B breast cancer showed a similar trend towards worse survival, the results did not reach statistical significance. As additional proofs-of-principle for the use of archetypes to predict overall patient survival, we identified archetype-associated groups with poorer survival in two additional TCGA cohorts: Colon Adenocarcinoma (*N* = 481) and Pancreatic Adenocarcinoma (*N* = 178) ([16, 17], cf. SI Fig. 2). In summary, clinical treatment of breast cancer is consistent with predictions based on their archetypes; however, the general applicability and effectiveness of archetypes in distinguishing patient survival outcomes suggests that they may offer superior, tumor-type agnostic capabilities for patient stratification.

Finally, to characterize archetype enrichment patterns in a representative metastasis model, we examined an additional dataset comprising median gene expression profiles of basal breast cancer from primary tumors and five distinct metastatic sites: brain, lung, liver, adrenal gland, and skin ([20, 26], cf. Methods). Consistent with [26], breast metastases exhibited archetype enrichment patterns resembling their destination organs more than their primary tumor site (cf. Fig. 4E). However, these distant metastases were less enriched for destination archetypes compared to primary tumors from the same location, indicating an intermediate behavior (cf. Fig. 4F). Nevertheless, these findings, in conjunction with our previous results, could suggest that metastatic acquire treatment sensitivities associated with their destination archetypes.

### DNA alterations guide, but do not predict, the onset of non-canonical tissue signatures

We next used CCLE to evaluate the association between tumor archetype scores, copy number alterations (CNA), and mutations (cf. Methods and Fig. 5). Our analysis revealed significant Spearman rank correlations (BenjaminiHochberg *p <* 0.05) involving 18 out of 73 TCGA considered hotspot mutations and at least one normal tissue archetype (cf. Fig. 5A). Notably, *TP53* mutation was one of the strongest correlates with the Proliferation archetype, aligning with the expected link between *TP53* dysregulation and a shift towards the most commonly enriched tumor archetype across various cancer cell lines (cf. Fig. 2A). However, the effect size was relatively modest, with *TP53* mutation leading to an average increase in the Proliferation archetype score of approximately 0.1 (cf. Fig. 5B).

**FIG. 5:**
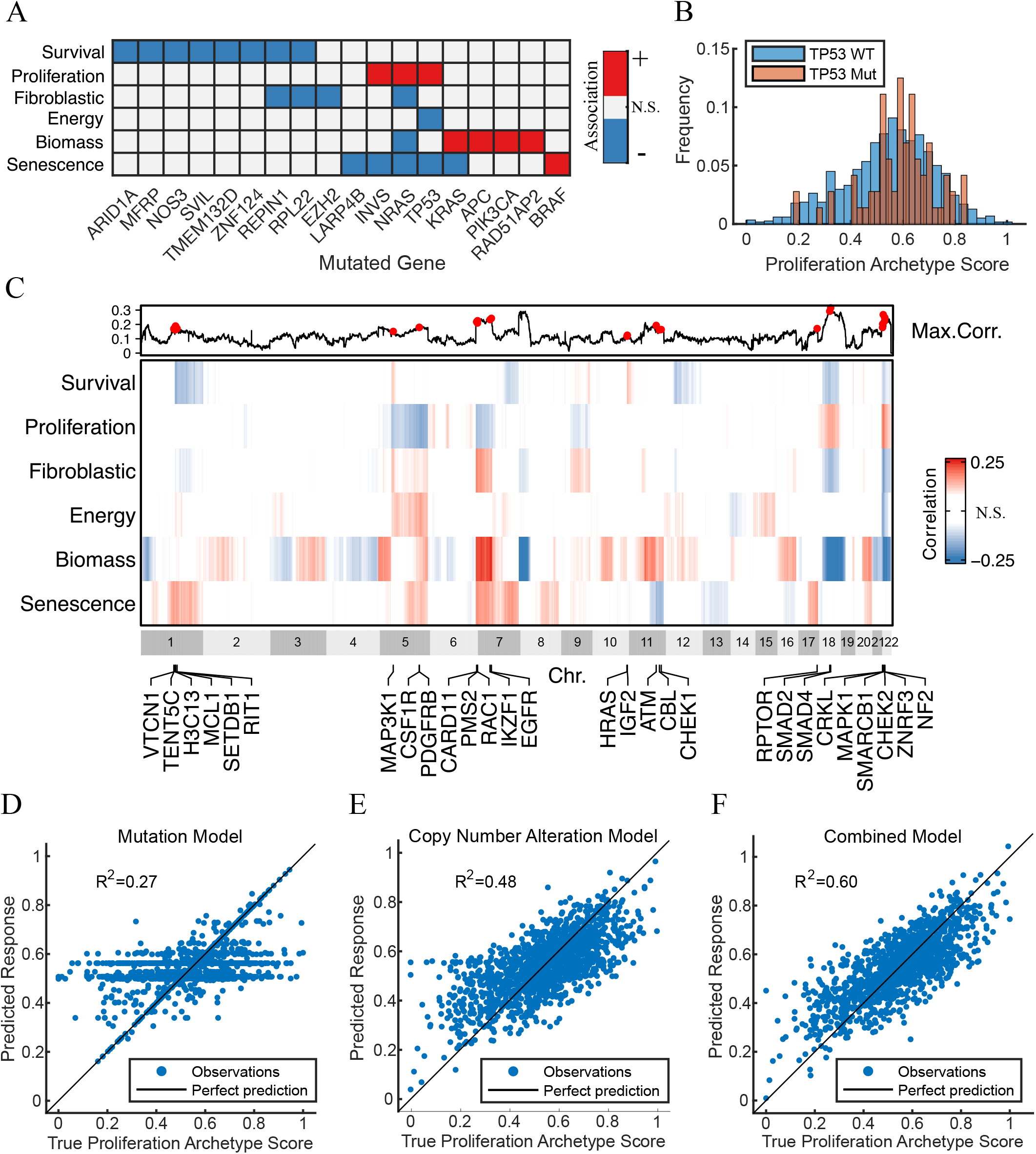
Associations between normal archetypes, copy number alterations (CNA), and mutations in cancer cell lines. (A). Heatmap of the significant associations between individual archetypes and recurrent TCGA hotspot mutations found in CCLE (Spearman rank permutation test, Benjamini-Hochberg *p <* 0.05). (B). Histograms of the Proliferation archetype score in cell lines with (orange) and without (blue) a TP53 mutation. (C). Heatmap of the significant genome-wide bin-averaged associations between individual archetypes and gene-level CNAs found in CCLE (Spearman rank permutation test, Benjamini-Hochberg *p <* 0.05). The inset above shows the maximum absolute correlation of each bin across the individual archetype scores. Genes labeled and marked in red on the inset were found in both in MSK-IMPACT and in the top 1% of correlations to one of the six archetypes. (D). Linear prediction of the Proliferation archetype scores from the 73 recurrent hotspot mutations retained in CCLE. (E). Linear prediction of the Proliferation archetype scores from the top 500 principal components of the gene-level CNA data in CCLE. (F). Linear prediction of the Proliferation archetype scores from the combined features used in D and E. All regressions were performed using the MATLAB Regression Learner Application. CNA regression analysis was restricted to genes from MSK-IMPACT.

Our investigation also identified several CNAs significantly associated with normal tissue archetype scores (cf. Fig. 5C). Each archetype was found to be associated with a distinct set of CNAs, similar to our findings regarding mutations. Moreover, multiple regions containing CNAs measured in MSK-IMPACT [48] were among the top 1% of Spearman correlations (cf. Fig. 5C, inset). This suggests that whole-genome DNA alteration profiles may be able to predict details about tumor archetypes.

To test this hypothesis, we conducted linear regressions using previously identified features, including mutations alone (Fig. 5D), CNAs from MSK-IMPACT alone (Fig. 5E), and mutations combined with CNAs (Fig. 5F). Despite incorporating all 73 hotspot mutations and the top 500 principal components of the CNA data, no regression yielded an *R*^2^ value exceeding 0.60. Consequently, our analyses suggest that normal tissue archetypes offer independent prognostic information about tumors beyond the predictive capacity of the considered genetic features.

## DISCUSSION

We introduced a strategy to standardize tumor phenotype quantification across diverse cancers and datasets by identifying conserved transcriptional signatures called archetypes. In doing so, our analysis identified six archetypes – Survival, Proliferation, Fibroblastic, Energy, Biomass, and Senescence – defining common modes of transcriptional variability in both normal tissues and multiple cancers. This universality allows the prediction of disease characteristics, such as drug sensitivities and overall patient survival, across multiple cancers using standardized signatures. Importantly, their association with normal tissue transcriptional modules suggests a connection between site-specific adaptive processes during metastasis and potential therapeutic vulnerabilities. These features, not predictable from mutations and copy number alterations alone, may enhance prognostic capabilities alongside established DNA-alteration-driven personalized frameworks in cancer medicine [49].

Site-specific archetype adaptation may present a novel therapeutic vulnerability in metastatic cancers. Consistent with our findings, prior studies indicate that metastatic tumors express genes associated with their target tissues [26, 50]. Thus, the dual role of archetypes as markers for such tissue-specific gene expression patterns and predictors of tumor drug sensitivities suggests that site-specific archetype enrichment patterns may highlight vulnerabilities in metastatic tumors to treatments associated with the archetypes of their destination tissues. For instance, breast metastases to the liver, regardless of primary HER2+ status, are anticipated to be responsive to *EGFR* inhibitors such as lapatinib, suggesting a potential untapped therapeutic vulnerability [51]. The question remains whether tumors adapt their archetypes post-metastasis or if cell populations enriched for destination-specific archetypes migrate to those locations [52]. If the latter, interventions could potentially steer primary tumor archetypes, guiding eventual metastases to more manageable sites [53].

Our findings suggest that DNA alterations and archetype enrichment offer distinct yet complementary information. Indeed, recent studies support the notion that DNA alterations alone do not reliably predict transcriptional behaviors or where a cancer will metastasize to [54, 55]. Nevertheless, our alteration correlation map (cf. Fig. 5) may aid in interpreting the impact of specific mutations and copy number alterations on tumor phenotype. Conversely, DNA alterations may act, in part, as archetype regulatory elements, steering tumors into specific behaviors and consequently modulating their drug sensitivities. For instance, our analysis associates mutation in *BRAF* with a shift towards the treatment-resistant Senescence archetype, aligning with its role in treatment-resistant cancers [14]. A noteworthy implication is that such alterations might inadvertently sensitize tumors to therapies associated with their new archetype compositions, a phenomenon termed collateral drug sensitivity [56], warranting further investigation in future studies.

The archetype mixture model was trained using normal tissues to maximize signal-to-noise and to compare them with known cellular trade-offs. Furthermore, while we restricted our gene set to five hallmark pathways of broad relevance to cancer, additional archetypes could be contained in the remaining genes. However, even with these restrictions, our archetypes are in broad agreement with known cancer hallmarks [8]. Furthermore, the utilization of N-NMF allows our framework to be easily augmented with additional archetypes and data constraints, allowing our foundation to be used as a starting point for additional development [28].

There are important caveats and considerations for the current analyses. First, the N-NMF method requires nonnegative data, rendering it unsuitable for application to z-scores or other transformations containing negative values. Second, the process of data normalization can potentially introduce spurious gene anticorrelations. Consequently, when generating archetypes through N-NMF, it is advisable to either compare them with clusters identified within the unnormalized data or with established biological labels, as we did when we mapped the archetypes to distinct tissues. Third, despite some evidence to the contrary [26], bulk tumor archetype enrichment patterns might be confounded by stromal tissue infiltration. Fourth, bulk tumor archetype scores significantly differed in scale from those in cancer cell lines used for drug sensitivity assessments. Therefore, additional investigation is needed to validate the present drug sensitivity predictions in bulk tumor samples.

In summary, six transcriptional signatures identified in normal tissues characterized heterogeneity patterns across a diverse range of human cancers. These signatures predicted both drug sensitivities and prognosis, going beyond what could be anticipated from DNA alterations alone [54, 57, 58]. Notably, these signatures are specifically concentrated in certain sites of basal breast cancer metastases, suggesting a potential new approach for treating metastatic cancers. Ultimately, recognizing the clinical significance of shared transcriptomic behaviors across human cancers may lead to complementary therapies targeting standardized signatures rather than cancer-specific features [53, 58].

## AUTHORS’ CONTRIBUTIONS

Conceptualization: C.W.; Methodology: C.W., K.A.M, and J.Z.; Analysis and investigation: C.W. and K.A.M.; Writing: C.W., K.A.M., and J.O.D.; Editing: C.W., K.A.M., J.Z., L.N., K.A.D., A.R.T, and J.O.D.; Supervision: A.R.T. and J.O.D.; Funding acquisition: C.W., L.N., A.R.T, and J.O.D.

## ACKNOWLEDGMENTS

Research presented here was funded by the following: the Marie-Josée Kravis Fellowship in Quantitative Biology (C.W.), the Laufer Center for Physical and Quantitative Biology (K.A.D.), AFOSR grants FA9550-20-1-0029 and FA9550-23-1-0096 (A.R.T.), NIH grant R01-AG048769 (A.R.T.), a grant from Breast Cancer Research Foundation BCRF-17-193 (J.O.D., L.N., and A.R.T.), Army Research Office grant W911NF2210292 (A.R.T.), and a grant from the Cure Alzheimer’s Foundation (A.R.T.).

