## Supplemental Figures for "Functional transcriptional signatures for tumor-type-agnostic phenotype prediction"

(Dated: March 12, 2024)

### SUPPLEMENTARY INFORMATION

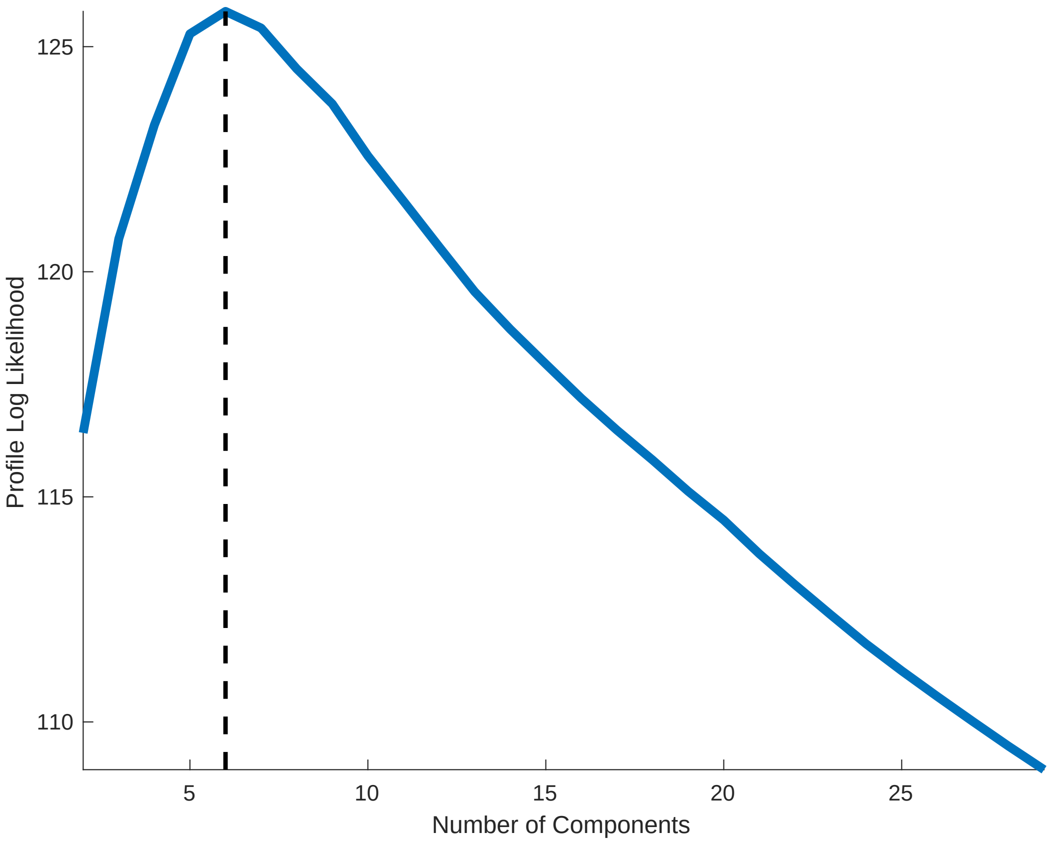


SI FIG. 1: **Profile log-likelihood analysis to determine optimal N-NMF rank** $k$. Profile log-likelihoods were computed over each $k$ number of components to optimize the N-NMF algorithm when approximating the GTEx dataset. The black dotted line represents the maximum profile log-likelihood at the optimal $k = 6$.

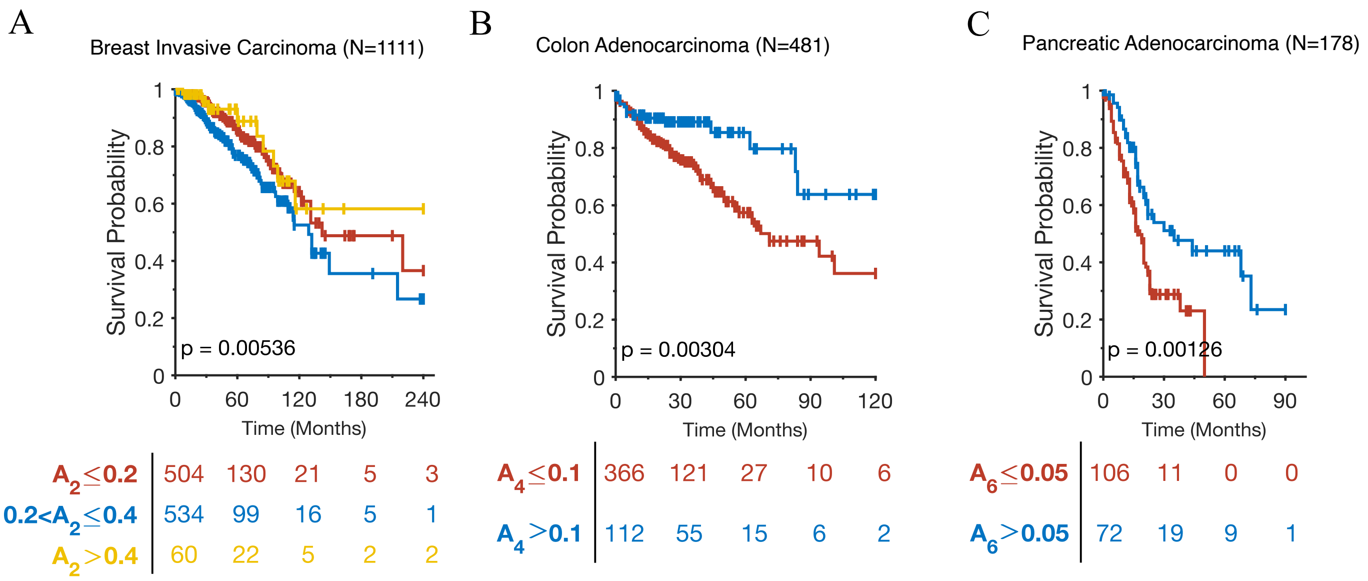


SI FIG. 2: **Normal tissue archetypes define groups with poor survival in (A) Breast, (B) Colon, and (C) Pancreatic cancers from TCGA.** Kaplan-Meier survival curves were stratified by archetype score. Tables below indicate the number of surviving patients at each time point. Significance was determined using a log-rank test. A is repeated from Fig. 4C to facilitate comparison.
